# Barrett’s esophagus is the precursor of all esophageal adenocarcinomas

**DOI:** 10.1101/2020.05.14.096826

**Authors:** Kit Curtius, Joel H. Rubenstein, Amitabh Chak, John M. Inadomi

**Affiliations:** Centre for Genomics and Computational Biology, Barts Cancer Institute, Queen Mary University of London, London, United Kingdom; Barrett’s Esophagus Program, Division of Gastroenterology, University of Michigan, Ann Arbor, Michigan, USA; Center for Clinical Management Research, Ann Arbor Veterans Affairs Medical Center, Ann Arbor, Michigan, USA; Division of Gastroenterology and Liver Diseases, Case Western Reserve University, Cleveland, Ohio, USA; Division of Gastroenterology, University of Washington, Seattle, Washington, USA

## Abstract

**Objective:** Barrett’s esophagus (BE) is a known precursor to esophageal adenocarcinoma (EAC) but current clinical data have not been consolidated to address whether BE is the origin of all incident EAC, which would reinforce evidence for BE screening efforts. We aimed to answer whether all expected prevalent BE, diagnosed and undiagnosed, could account for all incident EACs in the US cancer registry data.

**Design:** We used a multi-scale computational model of EAC that includes the evolutionary process from normal esophagus through BE in individuals from the US population. The model was previously calibrated to fit SEER cancer incidence curves. Here we also utilized age- and sex-specific US census data for numbers at-risk. The primary outcome for model validation was the expected number of EAC cases for a given calendar year. Secondary outcomes included the comparisons of resulting model-predicted prevalence of BE and BE-to-EAC progression to the observed prevalence and progression rates.

**Results:** The model estimated the total number of EAC cases in 2010 was 9,970 (95% CI 9,140 – 11,980), which recapitulates all EAC cases from population data. The model simultaneously predicted 8-9% BE prevalence in high-risk males age 45-55, and 0.1-0.2% non-dysplastic BE-to-EAC annual progression in males, consistent with clinical studies.

**Conclusion:** There are no additional EAC cases that plausibly arise in the US population outside the BE pathway. Effective screening of high-risk patients could capture the majority of population destined for EAC progression and decrease mortality through early detection and curative removal of small (pre)cancers during surveillance.

**Summary Box:** **What is already known about this subject?**

- Barrett’s esophagus (BE) patients have a 40 to 50-fold higher risk of developing esophageal adenocarcinoma (EAC) than the general population yet many remain undiagnosed.
- Identified BE patients receiving surveillance can have early cancers discovered endoscopically, which decreases the high overall EAC-associated mortality.
- Currently around 90% of patients who develop EAC were never part of a BE surveillance program, and those BE patients on surveillance have a low annual progression rate of 0.1 - 0.3% to develop EAC.

**What are the new findings?:**

- By applying a model that incorporates the evolution from normal cells to BE to EAC in patients, we found that the numbers add up - the expected number of EAC cases in the US population are explained by the published rates of BE described above.
- We cohesively examined the published estimates to determine that all EAC likely arises from both identified BE and occult, undiagnosed BE in the population.

**How might it impact on clinical practice in the foreseeable future?:**

- Based on current best estimates, our findings suggest there is no public health need to seek cases of a non-BE alternative pathway to EAC.
- Increasing efforts for effective, sensitive screening and surveillance of the true BE population will decrease EAC mortality in the coming years.

## Introduction

Esophageal adenocarcinoma (EAC) is typically diagnosed when a patient presents with symptoms such as dysphagia. Unfortunately, the majority of these patients do not live past the first year of their diagnosis because by the time dysphagia develops, metastatic cancer is already present. In order to prevent this cancer or detect it at an earlier, more treatable stage, efforts are now made to identify patients with Barrett’s esophagus (BE), the only known precursor to EAC. Identified BE patients are believed to have a **40 to 50-fold** higher annual incidence of EAC than the general population.^1^ Metaplastic BE progresses through dysplasia to cancer. Advances in endoscopic eradication therapy for dysplastic BE discovered during surveillance of BE can now prevent cancer.^2^ However, most cancers arise in patients without previously diagnosed BE suggesting either inadequate screening strategies or, as a recent study proposes, the possible existence of a pathway independent from the BE pathway.^3^ In this study we seek to answer a simple question about the unseen origins of EAC: **does overall EAC incidence reflect the number of cancers that would be expected to arise from prevalent BE?** In other words, do any EAC cases remain unaccounted for that ergo did not arise from the typical Barrett’s precursor pathway? The answer to this question will importantly guide research and public health efforts. If BE is the major or only precursor of EAC, then investigators should continue to focus on improving BE detection. If BE is not the major precursor of EAC then research needs to focus on identifying alternative pathways and BE screening programs will have limited impact on prevention and early detection of EAC.

In reality, very few individuals who have BE are ever offered an upper endoscopy, and therefore most BE remains asymptomatic and undiagnosed. Patients with gastroesophageal reflux disease (GERD) are technically the only sub-population of the general public typically recommended BE screening because it is believed they have a **5-fold relative risk** of developing long segment BE,^4^ yet even so only about **10% of GERD patients will receive an endoscopy.**^1^ This indicates *under-screening*, likely because patients either do not complain of their GERD symptoms, they respond adequately to medical therapy, or were otherwise not deemed suitably high-risk by their physician to warrant an esophagogastroduodenoscopy (EGD). Nonetheless, the **prevalence of BE in the general population is 1-2%**, whether diagnosed or not ^5,6^, and this is likely considerably higher in certain at-risk groups in the US.^7-10^ The main concern is that the average rate to develop EAC in these patients is low – around **0.3% per year.**^11^ Therefore, the majority of endoscopies are futile in finding EAC. We aimed to answer whether all prevalent BE we expect to find, diagnosed and undiagnosed in the US population, could account for all the incident EACs we expect to find as progression rates would imply, to fit the national cancer registry data.

## Methods

The question above is too complex to answer on the ‘back of an envelope’ because published *average* rates of progression are dependent on age, birth cohort, and calendar year. In particular for EAC, age-specific incidence rates vary drastically between men and women.^12^ This complexity of timescales involved in normal to premalignant BE to EAC progression has necessitated the creation of quantitative models that analyze cancer incidence rates, and project these trends into the future for public health risk assessments and planning.^13^ Models also quantify the potential impact of progression rates measured in clinical studies on hypothetical intervention and surveillance scheduling in efficacy and cost-effectiveness studies.^14-16^ Such models allow us to perform quantitative, comparative analyses on the benefits versus harms of proposed screening and surveillance protocols against watch-and-wait strategies; these simply cannot be done heuristically due to the complex nature of cancer evolution.

In this study we model both the onset of BE and the progression of BE to EAC. As a brief background, the multistage clonal expansion for EAC (MSCE-EAC) model is a stochastic model for development of EAC during patient lifetime that includes probabilities of developing BE at various ages, followed by initiation of dysplastic and malignant cell clones in BE with parameters for growth and progression of individual clones to cancer (Figure 1). The *inputs* only include GERD prevalence (calibrated to age- and sex-specific estimates^17, 18^) and EAC age- and sex-specific incidence curves provided by SEER registry.^12^ The BE prevalence and neoplastic progression rates are calibrated to fit those inputs, i.e. they are not based on observed BE prevalence or neoplastic progression rates from empiric studies. Briefly, the model includes GERD-stratified risk curve to develop BE, which is modeled as an age-dependent rate of exponential BE onset each calendar year with an unknown baseline parameter *ν*_0_. The patient-specific BE lengths can vary, derived from a Beta distribution with general population mean length set to 2 - 3 cm. Beyond *ν*_0_, the baseline constant rate for BE onset, the additional model parameters govern the evolutionary dynamics for dysplastic and malignant growth and EAC detection. The model parameters have been previously calibrated such that the resulting hazard functions fit to EAC age- and sex-specific incidence curves provided by SEER registry.^13^ We found during rigorous model selection with likelihood ratio tests that models stratified by birth cohort and sex best fit the incidence data, robust to sensitivity analyses (Figure 2). With these fits, the model *outputs* used for this study include the expected number of EAC cases in an at-risk population at a given year calculated using the hazard function *h*_*EAC*_ (see Supplementary material for equation details), along with the BE prevalence and the resulting BE-to-EAC progression rates (predicted as specific to age, sex, and birth cohort).

**Figure 1:**
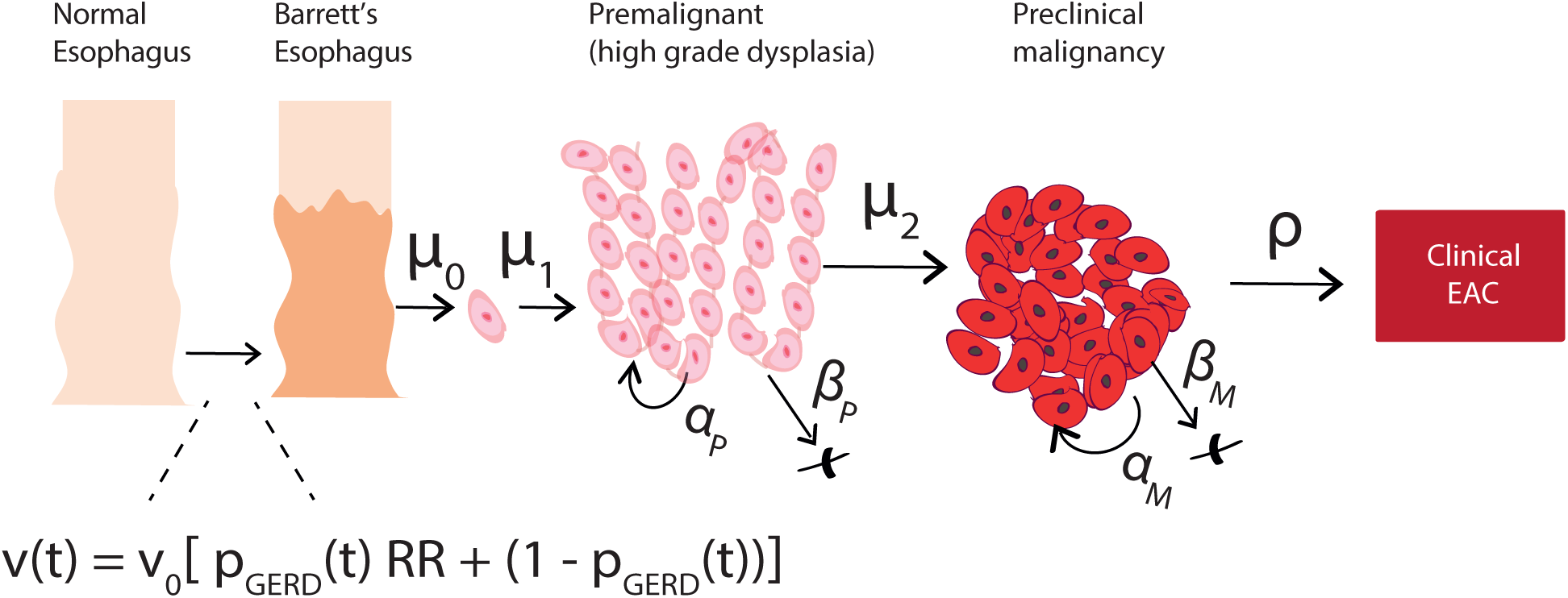
The stochastic, multi-scale model for EAC development (MSCE-EAC) includes conversion from normal squamous epithelium in the esophagus to BE metaplasia with BE onset rate *ν*(t), which is a function of a baseline rate *ν*_0_ and age-dependent prevalence of GERD p_GERD_(t) (see Methods for details). Two-hit processes with rates μ_0_, μ_1_ can initiate a premalignancy (e.g., inactivation of tumor suppressor gene *TP53* in non-dysplastic BE due to mutation/copy number alteration in a BE daughter cell creates first cell of a high grade dysplasia lesion). Premalignant cell growth rates are defined as α_P_ = division rate, β_P_ = death/differentiation rate per year. Malignant transformation with rate μ_2_ creates the first cell of a pre-clinical clone that can grow with rates α_M_ = division rate, β_M_ = death/differentiation rate per year. Size-based probability ρ for detection of preclinical malignant clone can lead to patient-specific time of incident EAC.

**Figure 2:**
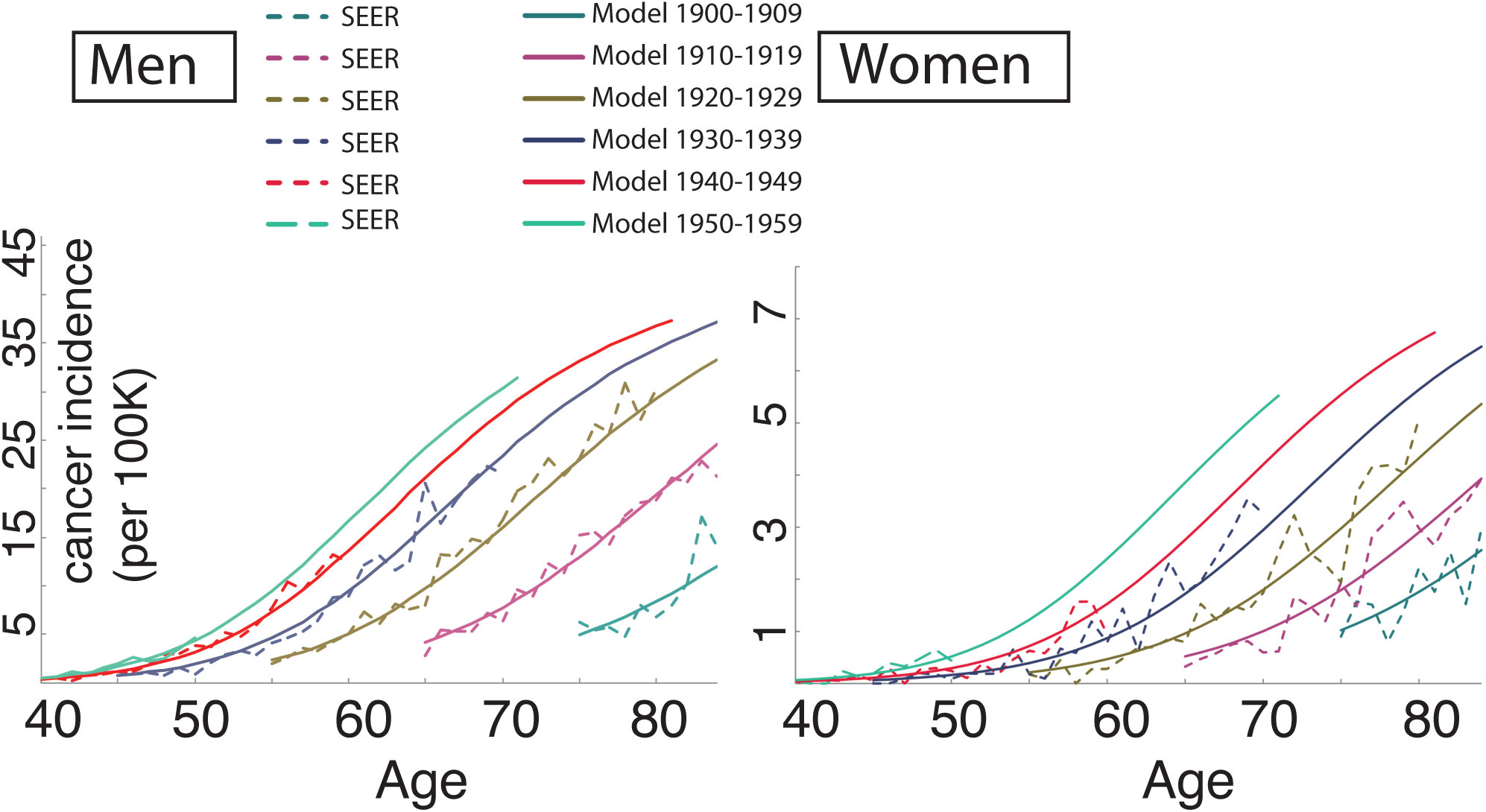
MSCE-EAC model was previously calibrated to EAC incidence curves (solid lines), stratified by sex and birth cohort, for consistency with SEER data trends (dashed lines).

This model has been used and improved in comparative analyses within the NCI Cancer Intervention and Surveillance Modeling Network (CISNET) consortium for the past 9 years, which has enabled numerous studies on sensitivity of biopsy sampling techniques for detection of small dysplastic lesions^14^, on influence of patient-specific molecular BE dwell time on future EAC risk^19^, and on cost-effectiveness of endoscopic eradication therapy for certain BE risk groups during surveillance.^15^ In our original study on modeling EAC incidence and mortality rates from 1975-2010, we used SEER-specific model fits combined with US census data to estimate past and predict future EAC-related deaths but did not include predicted EAC cases by calendar year when applied to US census data.^13^

In the Results below, we expand upon prior modeling to help elucidate an answer to our general public health question – “Is BE the precursor of all EAC?” To do this, we first applied the model to estimate the number of EAC cases using the US age- and sex-specific at-risk population estimates from the US census data, to be able to compare with the expected number quoted by Vaughan and Fitzgerald.^1^ This outcome serves as an independent validation of successful calibration of our model to EAC incidence. Then we compared the simultaneous predictions of age-specific BE prevalence using the MSCE-EAC model with the published data currently used for screening rationale,^20^ which included endoscopic reports from the Clinical Outcomes Research Initiative (CORI) for more than 150,000 patients, most of whom were born around 1950. We also compared the mathematical predictions of neoplastic progression rate from non-dysplastic Barrett’s esophagus to published estimates.

## Results

First, Vaughan and Fitzgerald estimated that the newly diagnosed number of cases for ages greater than 40 to be roughly around 10,000 total in the US every year based on data from 2010 with an average EAC incidence rate across all age groups.^1^ With the Markov model framework we can analytically compute the EAC hazard function and estimate the expected number of newly diagnosed EAC cases by age and year separately for men and women when considering also population data. As a starting point using 2010 census person-year data,^21^ the model predicts about **2.2 million adults had prevalent BE** in 2010, which is around **1.6% of the general US population** over age 40. Then, for age groups greater than 40 in both sexes of all races, our single-age calibrated model estimated that **the expected number of new EAC cases diagnosed in 2010 was equal to 9**,**970 (95% CI 9**,**140 – 11**,**980).**

We also computed the analogous estimate for EAC cases using incidence rates quoted directly from the SEER registry for ages 40-90, which was found to be 9,400 EAC cases total in 2010.^12^ Thus, the estimate generated by our computational model of progression from BE to EAC is consistent with the total number of EAC cases reported in SEER, which also aligns with the 10K incident cases quoted by Vaughn and Fitzgerald.^1^

Secondly, we considered what the model simultaneously predicted for BE prevalence and BE-to-EAC progression rates in order to achieve the expected ∼10K cases. Breaking down the contributions of the 2.2 million total BE patients estimated above, the model predicted **BE prevalence to be 1.9% - 2.4% in men and 0.4% - 0.5% in women in the general US population ages 45-55 in 2010** (Figure 3A). These predictions concur with best estimates^5, 6^ and influence the total EAC cases predicted by the multi-stage model. To further explore implications for high-risk patients, we note the model predicted a **BE prevalence of 7.9% - 9.3% in US men with symptomatic GERD who are cancer-free ages 45-55 in 2010** when the relative risk (RR) of BE versus non-GERD individuals is assumed to be RR=5 (Figure 3B). This is also consistent with the estimate of 8% provided by Vaughan and Fitzgerald ^1^ for prevalence of cancer-free BE diagnoses amongst GERD patients who undergo an upper endoscopy. Further, the model’s predicted age-specific BE prevalence curves by sex were consistent with previous results on BE prevalence from the Clinical Outcomes Research Initiative (CORI) study ^20^ (Figure 3A). Compared to our model results and 8% quoted above^1^ for high-risk groups, the CORI study independently found similar BE prevalence in white men with GERD of 6.3% for ages 40-49 and 9.3% for ages 50-59 (Figure 3B). To account for likely heterogenous relative risk of developing BE in GERD populations based on symptom onset age, BE length, and other factors,^4, 22-26^ we also considered a range of fixed values (RR = 2 – 6) and found age-specific trends broadly consistent to overall BE prevalence results in CORI. Observed BE prevalence in white women undergoing screening was less precise in the CORI study data yet still coincided with our predictions for women (Figure 3B).

**Figure 3:**
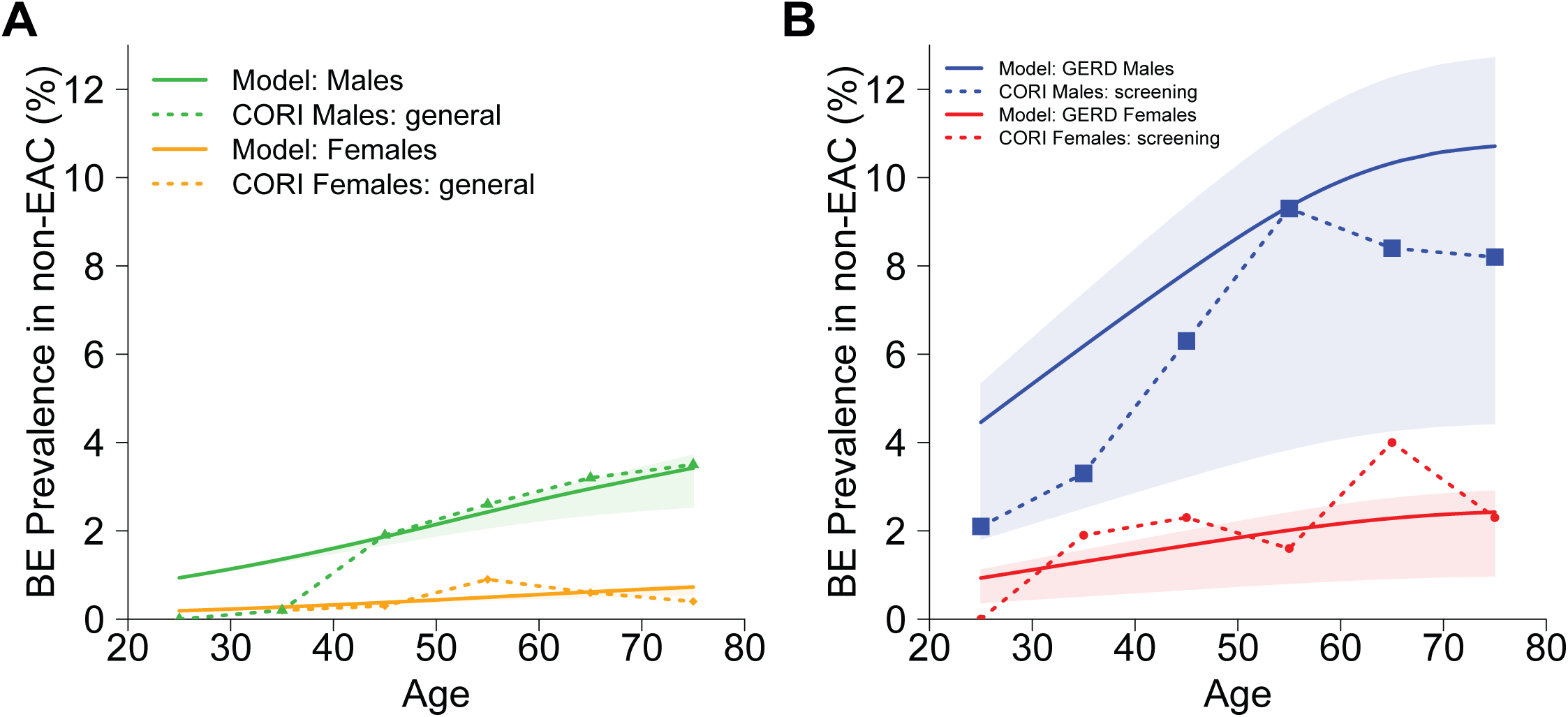
Model predictions for BE-positive yield in a cancer-free population (solid lines) are consistent with observed data (dashed lines) from Clinical Outcomes Research Initiative (CORI).^20^ **(A)** Solid lines show model results for the general US population stratified by sex from the 1950 birth cohort, with contributions of relative risk (RR) of BE from the age-specific, prevalent GERD population assumed to be RR=5 (shaded areas, RR=[2,6]). Dashed lines show consistency with observed BE prevalence data for patients without indication for screening in CORI, which are independent of the model. Model BE prevalence estimates are part of the evolutionary multi-stage process and thus affect predictions of the total EAC cases predicted (see Results). **(B)** Solid lines show model results for the symptomatic GERD subpopulation stratified by sex from the 1950 birth cohort with relative risk for BE set to RR=5. The shaded areas are predicted ranges for GERD subpopulations with fixed RR = 2 - 6 to describe a wide range of increased risks of BE in published estimates, based on factors such as onset age of GERD and BE length. The true GERD-specific BE prevalence contributing to mathematical formulation used in **(A)** is within this region, where individual contributions are based on GERD onset age and underlying distribution of RR. Dashed lines show BE prevalence data for patients with GERD, and/or another indication for screening, in CORI.

Lastly, we previously found using this model that, for individuals born after 1940, the range of **progression rates from BE-to-EAC was 0.10% to 0.20% for men**, and this was about twice as high as we found for women.^13^ These are plausibly low rates compared to current best estimates. ^11,27^ If there were a significant alternate non-BE pathway for EAC development, then this model (which does not include a non-BE pathway) would have estimated either a lower predicted population incidence of EAC than what was observed in SEER, a greater prevalence of BE than what was observed, or a greater rate of neoplastic progression among non-dysplastic BE than observed.

## Discussion

Based on the published epidemiology of BE and EAC, our analysis suggests that a major alternative non-BE pathway to EAC is an unlikely scenario. The existence of such an alternative pathway was suggested by a retrospective analysis of macroscopic reports of EAC specimens diagnosed without BE in two cohorts from the US and UK by Sawas and colleagues; however, their study conclusions remain speculative due to some important limitations including 1) a lack of longitudinally followed cases to EAC from non-BE patient esophageal tissue, and 2) the plausibility that small BE segments were completely overtaken by malignant expansions and thus were unmeasurable at cancer diagnosis.^3^ Moreover our result that BE is the main origin of EAC does not necessarily refute the existence of differing phenotypes for EAC - the finding that presence of BE was associated with better survival could plausibly be explained by the theory that more aggressive cancers are likely to replace the precursor BE more readily than less aggressive cancers.

Indeed, genetic and epigenetic analyses have also consistently shown BE and EAC to be very similar,^28-31^ and one study that sought genomic differences between adenocarcinomas with and without BE failed to reveal molecular differences between the two.^32^ Nonetheless, this is fortunate news that, with adequate uptake, screening for BE by upper endoscopy or minimally-invasive non-endoscopic technologies^16, 33^ could potentially identify and enroll all patients who are at risk for developing EAC into a surveillance program. Practically speaking, this would require very sensitive screening to detect all prevalent BE of any size and location (such as small BE patches at irregular Z-lines) but our results suggest these patients would constitute the majority of future EAC cases if detected.

Although the overall progression to EAC is low in patients diagnosed with BE, for those selected BE patients who have high grade dysplasia and/or early EAC detected during surveillance, effective treatment can save lives. In this way, our analysis reinforces the primary goal in BE screening for EAC prevention - that potentially all mortality caused by EAC in the general population could be prevented by effective surveillance of the entire BE population. Further, by mathematically analyzing the time-dependent nature of cumulative risk of BE in GERD patients, we can also use our multi-stage model framework to improve identification of at-risk populations by optimizing the timing of initial screening recommended for BE in symptomatic GERD.^34^ Although current intensive ‘one-size-fits-all’ surveillance strategies ^35-40^ would lead to high costs for those over-diagnosed BE screen cases and surveillance strategies clearly need to improve, we conclude that there is a strong rationale for screening for BE to reduce EAC mortality.

### Data sharing statement

All data used in our analysis are publicly available.^12,20^ All equations are provided either in Figures and Supplementary material or were previously published along with model parameters.^13,14^ Code to solve equations was developed in R (version 3.6.1). Scripts to repeat our analysis will be made available at: github.com/yosoykit/BEtoEAC_Results.

## Acknowledgments

The authors thank the Clinical Outcomes Research Initiative (CORI) study for access to endoscopy data. This work was supported by funding from NIH grants CISNET U01 CA152926, U01 CA199336 (JMI, KC, JHR), K24 DK080941 (JMI, KC), and a UKRI Rutherford Fund Fellowship (KC). JHR is also supported by NIH grant U54 CA163059 (BETRNET). AC is supported by NIH grants U54 CA163060 and P50 CA150964.

