## Supplementary material for "Barrett’s esophagus is the precursor of all esophageal adenocarcinomas"

*Model hazard function and expected number of esophageal adenocarcinoma (EAC) cases*

We compared the model's predictions to the US population estimates for Barrett's esophagus (BE) and EAC quoted by Vaughan and Fitzgerald,[1] by combining age- and sex-specific model hazard rates with at-risk population estimates from US census data. [2] For expected EAC incidence, using the Markov model framework (see Figure 1) we can analytically compute the EAC hazard function $h_{EAC}$ (see Curtius et al. for full derivation with age-specific GERD-dependent BE rates incorporated explicitly [3]) and estimate the expected number of newly diagnosed EAC cases by age and year separately for men and women, using population data for the at-risk population numbers.[2] This is computed as:

$$\Lambda_{i,j} = PY_{i,j} h_{EAC}(a_i, b_k),$$

where $PY_{i,j}$ is the number of person-years at-risk in period $c_j$ of age $a_i$ and birth cohort $b_k = c_j - a_i$. For the Age-Cohort model, the birth cohort specific hazard $h_{EAC}$ (previously fit using person-year data for specified US populations directly from SEER [4]) can be written as $h_{EAC}(a_i, b_k) = h_{EAC}(t \mid t = a_i, b_k)$. For the main Results of incident EAC cases in $c_j = 2010$ and ages $a_i$ between 40-90, this was computed separately for men and women and the 95% confidence interval for this estimate of summed total cases was computed by re-sampling Markov Chain Monte Carlo posterior distributions of birth year- and sex-specific model parameter estimates[3, 4] for 100K bootstrap iterations. The equation below calculates the expected total number of EAC cases diagnosed in 2010, $\Lambda_{2010}$, which was equal to,

$$\sum_{i=40}^{90} \Lambda_{male_{i,2010}} + \Lambda_{female_{i,2010}} = \mathbf{9,970} \;\; [95\% \text{ CI}: \;\; 9,140 - 11,980]$$

For the analogous calculation using SEER incidence rates extracted from SEER*Explorer [5] for ages 40-90 in 2010 and census person-year data[2] we estimated 9,400 EAC cases total.

*Patient and Public Involvement*

We did not directly include PPI in this study, but the SEER database used here is updated by an NCI committee that seeks quality improvement with patient/public feedback welcomed.
